# Dual-system free-operant avoidance: extension of a theory

**DOI:** 10.1101/2023.05.24.542134

**Authors:** Omar D. Perez, Anthony Dickinson

## Abstract

Our theory of positively reinforced free-operant behavior (Perez & Dickinson, 2020) assumes that responding is controlled by 2 systems. One system is sensitive to the correlation between response and reinforcement rates and controls goal-directed behavior, whereas a habitual system learns by reward prediction error. We present an extension of this theory to the aversive domain that explains why free-operant avoidance responding increases with both the experienced rate of negative reinforcement and the difference between this rate and that programmed by the avoidance schedule. The theory also assumes that the habitual component is reinforced by the acquisition of aversive inhibitory properties by the feedback stimuli generated by responding, which then act as safety signals that reinforce habit performance. Our analysis suggests that the distinction between habitual and goal-directed control of rewarded behavior can also be applied to the aversive domain.

---

Sidman (1953) introduced free-operant avoidance by scheduling periodic shocks that rats could postpone or omit by pressing a lever. This form of avoidance has always been problematic for a contiguity-based reinforcement account because a rat will respond for many minutes in the absence of any external stimulus change to provide a source of reinforcement. However, a potential contiguous source of reinforcement was identified by Konorski and Miller in their 1936 report of one of the initial experimental studies of avoidance (Konorski, 1948 p. 228-232; 1967 p. 380-383). They initially established a Pavlovian defensive or aversive salivary conditioned response to a noise by pairing this signal with the delivery of dilute distasteful hydrochloric acid into the dog’s mouth. Once the salivary conditioned response was established, they occasionally passively flexed one of the dogs forelegs for 5 s during the noise and omitted the acid outcome. In the protocol illustrated by Konorski (1967 p. 381), after 7 noise presentations with the passive flexion, the dog for the first time spontaneously (and presumably voluntarily) flexed its leg during the noise, resulting in the omission of the acid delivery. Thereafter the dog flexed its leg many times during almost every noise presentation, thereby avoiding most of the impending acid deliveries.

Critically, Konorski and Miller also observed that the spontaneous leg flexions were accompanied by a marked reduction in the conditioned salivary response to the noise, which led them to suggest that feedback stimuli generated by the leg flexion had become conditioned aversive inhibitors through Pavlovian conditioning because these stimuli predicted the omission of an expected aversive outcome, the acid. Moreover, they argued that this property of the feedback stimuli instrumentally reinforced avoidance responding. We shall refer to this account of avoidance as the safety signal theory, which was the first instantiation of what has come to be called a two-process theory (Rescorla & Solomon, 1967) and has received more contemporary endorsement (Dinsmoor, 2001).

Although Konorski and Miller’s observations are compatible with the safety signal account, they did not experimentally manipulate the properties of the feedback stimuli to evaluate the theory. The first to do so was Rescorla, who distinguished between two components of the safety signal account. The first is that the feedback stimulus becomes an aversive inhibitor on a free-operant schedule. To evaluate this claim, Rescorla trained dogs on a free-operant avoidance schedule under which each shuttle response produced a brief auditory feedback stimulus before presenting this stimulus independently of responding to assess its conditioned properties. Relative to a control condition in which the stimulus had been presented randomly while the dogs were responding, the feedback stimulus suppressed avoidance, thereby demonstrating its inhibitory properties (Rescorla, 1968). Rescorla and LoLordo had previously demonstrated that an independently established Pavlovan aversive inhibitor would suppress the free-operant shuttling avoidance response (Rescorla & Lolordo, 1965).

The second component is that an aversive inhibitor acts as a positive reinforcer for the avoidance response. To address this issue, Rescorla initially trained his dogs on a concurrent schedule of a free-operant avoidance in which pressing either of two panels postponed the next shock (Rescorla, 1969). In the second stage the panels were removed and the dogs were given a Pavlovian inhibitory conditioning. The avoidance schedule was reinstated with the condition inhibitor from the prior stage being presented following each press of one of the panels. During this test stage the dogs showed a clear preference for the response producing the aversive inhibitor, thereby establishing the aversive inhibitor as a positive reinforcer. Weisman and Litner reached the same conclusion using bidirectional instrumental control assessment of free-operant avoidance in rats (Weisman & Litner, 1969).

The safety signal theory assumes that free-operant avoidance, although operationally an example of negative reinforcement with the shock as the reinforcer, is in fact functionally an example of positive reinforcement by the feedback stimuli generated by the avoidance response. This form of positive reinforcement was explained by Dickinson and Dearing in terms of an opponent process between appetitive and aversive motivational systems under which a conditioned aversive inhibitor activates the appetitive motivational system and thereby functions like a conditioned appetitive excitor (Dickinson & Dearing, 1979). A variety of evidence supports this functional equivalence – for example, appetitive conditioning is blocked when conducted in the presence of an aversive inhibitor (Dickinson & Dearing, 1979; Laurent, Balleine, & Westbrook, 2018).

## Habitual Avoidance

One popular approach suggests that positively reinforced instrumental behavior comes in two forms (Daw & O’Doherty, 2013; Dickinson, 1985; Dickinson & Perez, 2018; Dolan & Dayan, 2013): as habitual responses and as goal-directed actions. Experience with an instrumental contingency is assumed to strengthen habitual responding without encoding information about the reinforcer or outcome of the response. A classic example of such a mechanism is Thorndike’s law of effect according to which an association between a current stimulus and a response is strengthened when the response is followed by an effective reward (Thorndike, 1911).

By contrast, goal-directed behavior is mediated by a rational interaction between knowledge of the causal relation between the action or response and the outcome and the current value of the outcome, which can vary with the relevant motivational variables (Heyes & Dickinson, 1990). Such an interaction is goal-directed in the sense it is *directed* by knowledge of the action-outcome contingency and motivated by the representation of an outcome as a *goal*. Consequently, a goal-directed account of safety signal learning involves encoding a representation of the outcome, in this case feedback stimulus, as a goal of the instrumental action.

The canonical assay for distinguishing between habitual and goal-directed control is the reinforcer or outcome revaluation test (Adams & Dickinson, 1981). The rationale for this test can be illustrated by a revaluation test conducted by Fernando and colleagues to determine whether free-operant avoidance by rats is goal-directed or habitual with respect to a feedback stimulus that functioned as a safety signal (Fernando, Urcelay, Mar, Dickinson, & Robbins, 2014b). Their rats were trained on a free-operant variable cycle (VC) schedule, which consisted of a variable avoidance period followed by a shock period in which three foot-shocks were presented with a short interval between them before the next component was presented. A lever press during either avoidance or shock period terminated the current cycle so that any further programmed shocks in that cycle were omitted. Therefore, by pressing in each avoidance period the rat could avoid all of the shocks that were scheduled to occur after the variable avoidance period.

Furthermore, each lever press produced a 5-s auditory feedback stimulus, which was assumed to function like the endogenous feedback stimuli produced by pressing the lever and thereby enhance the salience of the sensory feedback produced by the instrumental response. In accord with the safety signal theory, in separate experiments Fernando and colleagues not only replicated Rescorla’s finding of a preference for an avoidance response that produced the feedback stimulus but also that the feedback stimulus contingency enhanced the rate of the avoidance response (Dinsmoor & Sears, 1973), as well inhibiting avoidance responding in its presence.

Following this training, the lever was withdrawn and the rats received non-contingent exposures to the feedback stimulus under morphine in the revaluation group, whereas the control group received the same exposure to the morphine and feedback stimulus but in separate sessions. In a prior experiment, the same authors had demonstrated that this treatment enhanced the reinforcing effect of the feedback stimulus on avoidance responding relative to the control group when tested in the absence of the shock. In the critical experiment, however, they tested the effect of the revaluation treatment in the absence of both the feedback stimulus and the shock. If the revaluation treatment acts by enhancing the capacity of the feedback stimulus to reinforce habitual responding, such an enhancement should not be observed in the absence of this stimulus during the extinction test. In contrast, if the feedback stimulus acts as a goal through a representation of its current value, the revaluation should enhance responding even during the extinction test. Critically, Fernando and colleagues failed to detect any effect of the morphine revaluation on extinction relative to the unpaired control group, suggesting that the safety signal operates through habit-based reinforcement rather than enhancing the value of the signal as a goal. As it stands, this inference is based on a null result in the extinction test and therefore the habit-based interpretation also requires the demonstration of an interaction between the revaluation effect and the type of test - extinction versus a reinforced test with response-dependent feedback stimulus. A subsequent reinforced test replicated the enhancement of responding observed in the previous experiment and yielded a significant interaction. In conclusion, the Fernando et al. (2014b) experiments suggest that the feedback stimulus acts as a positive reinforcer of habit learning and not by establishing a representation of the causal relation between the action and its feedback stimulus that is necessary for goal-directed control. By extension, we should expect that intrinsically generated feedback stimuli also function as such positive reinforcers.

## Goal-directed Avoidance

As well as investigating the nature of the avoidance maintained by feedback stimuli, Fernando and colleagues also used the outcome revaluation procedure to determine whether the current value of the shock plays a direct role in controlling free-operant avoidance (Fernando, Urcelay, Mar, Dickinson, & Robbins,2014a). Rather than revaluing the feedback stimulus as in their previous study, they sought to revalue the shock. To this end, they trained the rats on their VC avoidance schedule and then, in the absence of the lever, exposed their rats to non-contingent presentations of the shock under morphine in an attempt to reduce its aversiveness. If avoidance was motivated by the negative value of the shock interacting with knowledge of the negative causal relationship between responding and the shock, this treatment should have reduced responding. This reduction is exactly what they observed in an extinction test without the shock relative to a control group that had received the shock and morphine in separate sessions during revaluation.^1^ The reduced avoidance persisted during initial exposure to the avoidance schedule in a subsequent reinforced test until the aversiveness of the shock was re-established by experiencing it in the absence of the morphine. Taken together, this pattern of reduced avoidance following shock revaluation would be expected if lever pressing on their schedule was goal-directed with respect to avoiding the shock.

What is less clear, however, is the nature of the representations and processes underlying this goal-directed avoidance. Shortly after Rescorla reported evidence for a role of safety signals in avoidance, Seligman and Johnson (1973) published a seminal chapter in which, having critically reviewed classic two-process or -factor theories of avoidance (Rescorla & Solomon, 1967), they argued for a goal-directed account of avoidance in terms of Tolmanian expectations and preferences (Tolman, 1948). Specifically, they suggested responding is generated by the interaction of expectations of particular outcomes following different actions, the avoidance response and non-avoidance response, and the preferences among their outcomes, no shock and shock, respectively, an idea that has been endorsed more recently by Lovibond (2006). Another contemporary framework for analyzing goal-directed behavior is that provided by computational reinforcement learning (RL) (Sutton & Barto, 2018). For example, Wang and colleagues (2018) developed a model-based RL account of goal-directed avoidance, which argues that the agent learns stimulus state-action-outcome state transitions using state prediction-errors that are then deployed in the selection of the action that yields the preferred outcome state. Such state transition learning maps paradigmatically onto discrete-trial avoidance learning in which performing the avoidance response in the state generated by the presence of a warning signal for an aversive outcome leads to a state in which the outcome is omitted.

Although expectancy and model-based RL theories may provide accounts of discrete-trial goal-directed avoidance, at least in the case of human avoidance (Gillan, Morein-Zamir, et al, 2013), it is not clear that such theories can be applied to free-operant avoidance which involves neither a warning signal nor an explicit trial structure. However, the classic Sidman schedule arranges a fixed shock-shock (SS) and response-shock (RS) intervals which allowed Anger to argue that a free-operant response is followed by an internal transition generated by the response feedback stimuli (Anger, 1963). He argued that the state generated at long post-response interval is more aversive than that produced by short post-response intervals because the long-intervals, but not the short-interval ones, have been paired with the shock. As a consequence, an avoidance response after a long-interval produces a transition to a less aversive state, which could act as a goal. There are, however, two problems with applying this account to a VC schedule which is the only schedule that has been shown to support goal-directed avoidance. First, as the VC schedule does not involve fixed RS intervals and so it is not clear that responding on this schedule would produce the relevant state transition. Second, as we have already described, the Fernando et al. (2014b) study suggests that VC avoidance responses generate a feedback stimulus state that acts through habitual learning.

In the absence of a computational theory of free-operant avoidance, we have explored whether a dual-system theory that we developed to explain positively reinforced free-operant behavior (Perez & Dickinson, 2020) can also capture free-operant avoidance.

### Rate-correlation System

Within our theory (Perez & Dickinson, 2020), the represented strength of the action-reinforcer relationship acquired through instrumental training is referred to as the goal-directed strength, g, which is a function of the correlation experienced between rate of responding and rate of shock *r*. The propensity to perform the action determined by the product of *g* and the incentive value of the reinforcer, *I*, which we shall refer to as the response strength. For a rewarded action, both *g* and *I* are negative and so is its response strength, thereby engendering its performance. By contrast, in the case of avoidance both *g* and *I* are negative - the former because the action-reinforcer experienced correlation is negative and the latter because the reinforcer is aversive. So once again the response strength, and hence the propensity of performing an avoidance action, is positive. The outcome revaluation procedure employed by Fernando and colleagues should have decreased the negativity of the incentive value of the shock, thereby reducing the response strength and therefore avoidance responding in the absence of the shock.

Computationally, we suggested that *r* (and therefore *g*) is determined by a mnemonic system that deploys a short-term memory (STM) to compute the current local correlation between the rate of responding and the rate of reinforcement. As illustrated in Figure 1a, the contents of the STM consist of a number of time samples each of which records the number of actions and the number of outcomes that occur in that sample. At the end of each time sample, the current correlation between the response and outcome rates, *r,* is calculated across the time samples currently in the STM before one of the samples is randomly deleted from memory and the registration of responses and outcomes in a new sample is started^2^. Figure 1a illustrates the operation of a simplified mnemonic system with a STM with the number of time samples (N) set to 4 across three cycles of the memory, whereas Figure 1b displays a simulation of three cycles of a larger memory size (*N* = 10) to calculate the current experienced *r* at each memory recycle, which is then used to update the running average. Note how each memory cycle differs from the previous one in only one data point; the agent forgets one sample from the previous cycle while it adds a new sample. Responding in the new sample is determined by the strength of the experienced rate correlation, which yields the strength of goal-directed control (*g*).

**Figure 1.**
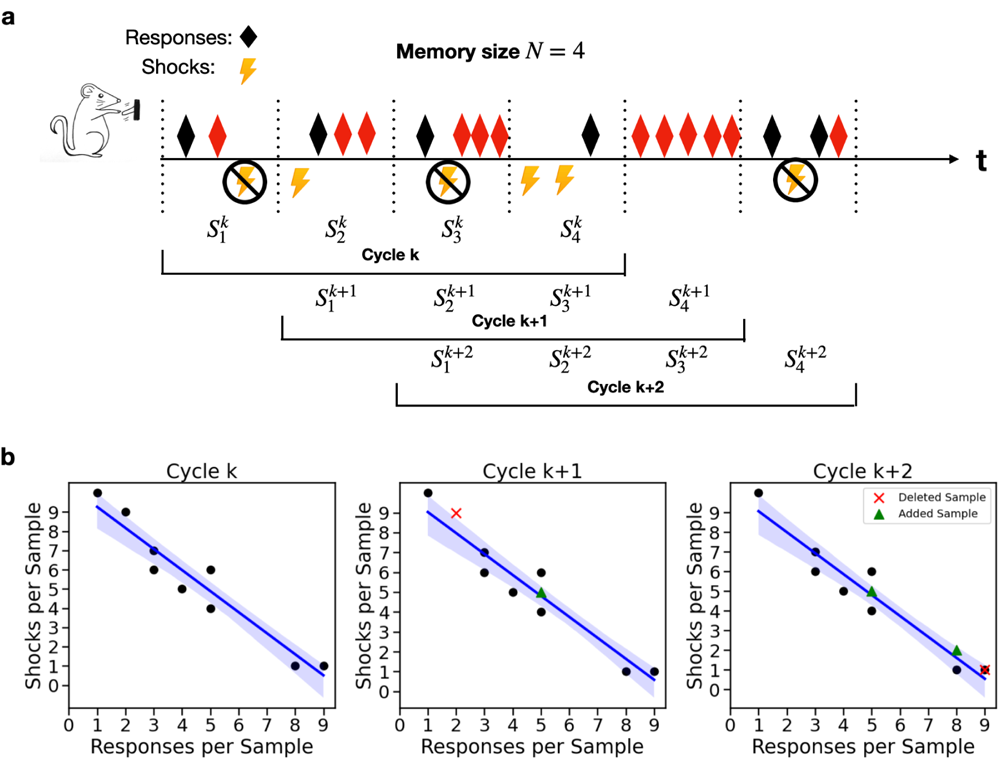
Avoidance free-operant training. a. Schematic representation of memory cycles in free-operant avoidance. In free-operant avoidance shocks are predetermined to come on a variable cycle (VC) schedule where a mean shock-shock (SS) interval is programmed by the experimenter using a random process. Subjects can respond at any time during a cycle. When a response is performed (black rhomboids in the figure), the next scheduled shock is canceled, but further responses within the cycle (before the shock is indeed canceled; red rhomboids in the figure) have no consequences on future programmed shocks. Each time-window represents a sample in short term memory (STM, with size N=4 samples in this simplified illustration). For each memory cycle *k*, the agent registers the responses per sample and received shocks per sample. After a memory recycle, one of the samples is randomly erased from memory (the first one in this example) and a new memory sample is added in the next cycle. **b. Simulation of memory cycles and their experienced rate correlation.** Each plot shows simulations of one memory cycle of size *N* = 10. In our rate-correlation system, events can be plotted as data points representing responses and shocks per memory sample. The blue line represents the best fitting line for each cycle; their slope represents the computed rate correlation experienced by the agent. One datapoint from memory cycle *k* (red cross) is randomly erased from memory while another is added for the next cycle *k* + 1 (green triangle); the same logic applies for all future memory cycles during training. The negative value of *g* is multiplied by the negative incentive value of the aversive shock (*I*) to yield a positive response strength *Ig* from this goal-directed system.

The strength of goal-direct control, *g,* is a weighted mixture of the current *r* and the current mean *r*, which when multiplied by the current incentive value of the outcome (*I)* determines the strength of responding in the next time sample. Of course, the current *r* can only be calculated if the current STM has registered at least one response and one outcome. In other words, the agent will compute the rate correlation only if in any particular memory recycle there is at least one response and outcome registered in memory. In the absence of a registered response and/or outcome, *g* is determined by the mean *r* prior to the recycle, which is not then updated until current memory registers at least one of each event.

With the outcome incentive value set to one (*I* = 1), this goal-directed system yields qualitative matches to the rates of lever pressing by rats under ratio and interval schedules of positive reinforcement across variations in the probability of the reward per press, the reward rate, and the delay between a reinforced press and reward delivery using a consistent set of parameters (Perez & Dickinson, 2020). Of course, under a negative reinforcement, or an avoidance contingency, the rate correlation will be negative, and at issue is whether the same system can also account for the impact of variations in the important determinants of free-operant avoidance when the incentive value of the negative reinforcer is set to minus one (*I* =− 1) to reflect its aversive properties. As a consequence, the product of *g* and *I* yields a positive response strength, which in combination with the habit system determines the rate of responding.

### Simulations

To implement simulations of the contribution of the goal-directed system to avoidance, the STM consisted of thirty 20-s samples, which is the sample duration used for the simulations of positively reinforced behavior reported by Perez and Dickinson (2020). Simulations were performed using the R programming language and Python 3.

The response-reinforcer relationship was computed and assessed at each memory recycle by a correlation coefficient between the number of presses and shocks per sample across the current samples in the STM (see Figure 1b). The strength of goal-directed control, *g*, throughout the next memory sample *k* + 1, *g* , was set at a *k*+1 weighted mean of the correlation yielded by that last recycle and the mean of the correlations at all previous recycles, that is:

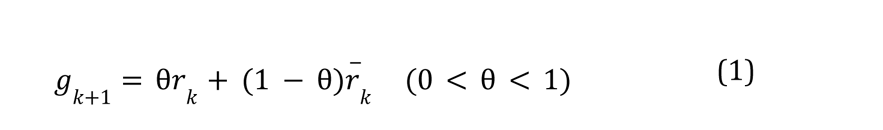

where *r*^-^_*k*_ is the mean correlation experienced during the experiment, up to memory recycle *k*. The weighting parameter θ controls the extent to which goal-directed control depends upon the most recent or distant experiences of the instrumental contingency between the action and the reinforcer, in this case the absence of a shock.

We assume that the probability of a response being performed at each second is constant in each memory recycle *k*, so that *p* = *Ig* ∼ *geom(p_k_*), where *geom()* is a *k k k* geometric distribution with parameter *p* , and *I* =− 1. Our initial simulations investigated the acquisition on a simple VC schedule studied by deVilliers (1974) that maintained a constant shock probability per second generated in a similar way to responding by using a geometric distribution with parameter *p* = *geom*(^1/T^), where *T* denotes the average interval between shocks, or the average SS interval. For example, for a VC 15-s schedule, shocks were generated by *s_t_ geom*(1/15) so that on average the SS interval was 15 s. Each response canceled the next programmed shock, and additional presses during the SS interval had no effect on subsequent programmed shocks. The parameter *T* (the mean SS interval) was varied across simulations to generate the VC schedules: 15 s, 30 s, 45 s, and 60 s. The weighting θ given to the current correlation relative to the mean correlation in determining *g* was set to 0. 8. *k*

The weighting parameter β which updates the rate correlation experienced was set to 0.5. In each of our simulations we report the mean response strength (*Ig*) across 200 *k* virtual rats, each consisting of 200 mnemonic recycles, at each schedule parameter (T) with an initial baseline press probability, *p_initial_*, set at .05 per second.

Figure 2a illustrates that the acquisition of response strength is relatively rapid with the terminal strength increasing with programmed shock rate as determined by the mean SS interval of the VC schedule. Bolles and Popp observed relatively rapid acquisition of lever pressing by rats on a standard Sidman avoidance once any interference from freezing was overcome (Bolles & Popp, 1964). Given that the simulated response strength rapidly becomes stable, even if variable, in all subsequent simulations we averaged the strength across the last 50 cycles. Figure 2b illustrates the impact of varying SS interval on the simulated goal directed strength corresponds to the profile of response rates observed by de Villiers (1974). In the next sections we assess whether the rate-correlation system can account for the two primary determinants of free-operant avoidance, shock rate reduction and experienced shock rate.

**Figure 2.**
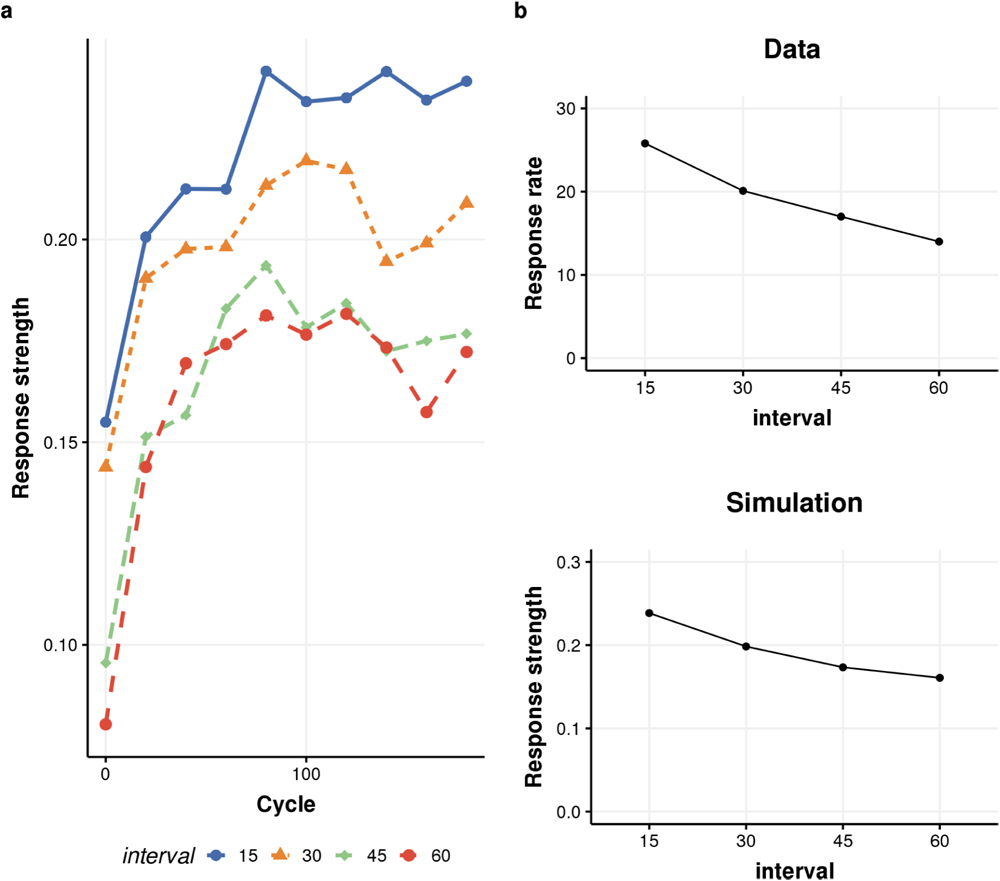
**a.** Mean response strength (*Ig*) across 200 memory cycles of the VC using the mean SS intervals studied by De Villiers (1974). **b. Top panel.** Mean responses per minute observed by de Villiers (1974). **Bottom panel.** Mean simulated response strengths (*Ig*) by the rate-correlation system.

### Shock Rate Reduction

Sidman (1962) himself suggested that free-operant avoidance is reinforced by the reduction in shock rate produced by responding, an idea that led Herrnstein and Hineline (1966) to investigate the role of shock rate reduction by varying the probability of a shock following a shock *p(shock|shock)* and following a response *p(shock|shock)*. When *p(shock|shock)* is lower than *p(shock|shock)*, responding reduces the shock rate until the next shock is received when the rate produced by *p(shock|shock)* is reinstated, a reduction that should have reinforced and maintained responding. In one analysis *p(shock|shock)* was set at .15 while *p(shock|shock)* varied between .05 and .15. As can be seen in Figure 3a, responding was maintained as long as *p(shock|shock)* was greater than *p(shock|shock)* but not when two probabilities were equal. The corresponding relationship was also observed in a second analysis when *p(shock|shock)* was fixed at .05 and the *p(shock|shock)* varied between .05 and .25^3^. As can be seen from Figure 3b and Figure 3c, the simulated response strength (*Ig*) values reproduce the basic functions observed by Herrnstein and Hineline, demonstrating that the rate correlation system in our model can capture the role of shock rate reduction in responding and goal-directed control.

**Figure 3.**
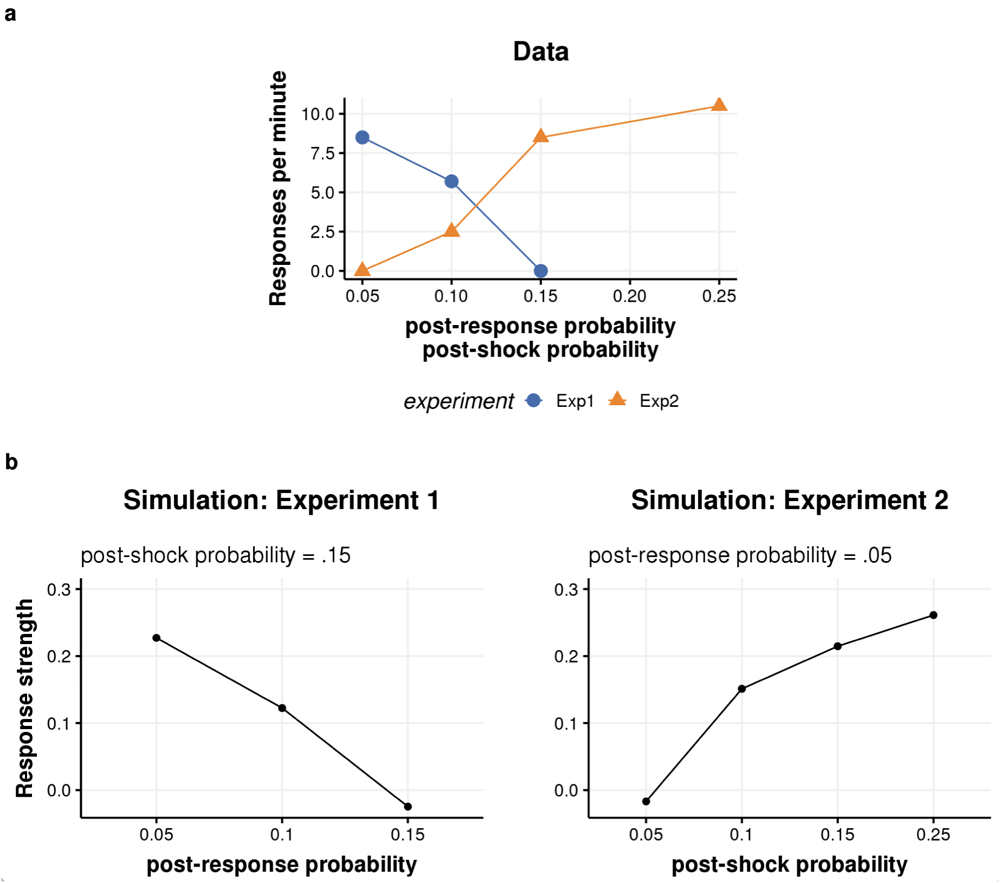
Simulation of shock rate reduction and response strength. **a.** Results obtained by Herrnstein and Hineline (1966). In Experiment 1 (Exp1, blue line) the probability of shock after having received a shock (the post-shock probability) was kept constant at .15 while the probability of shock after having performed an avoidance response (post-response probability) was varied as indicated in the abscisa. In Experiment 2 (Exp2, orange line), the post-response probability was kept constant at .05 and the post-shock probability was varied as indicated in the abscisa. **b & c.** Mean response strengths (*Ig*) obtained by a rate-correlation system for Experiments 1 and 2 by Herrnstein and Hineline (1966).

Responding by de Villiers’ (1974) rats on his simple VC schedule was not only systematically related to the scheduled SS interval (see Figure 2 Panel B) but also to the reduction in shock rate produced under each interval. As the left-hand panels of Figure 4 show, the response rate increased systematically with the reduction shock rate , a relationship reproduced by the simulations of response strength.

**Figure 4.**
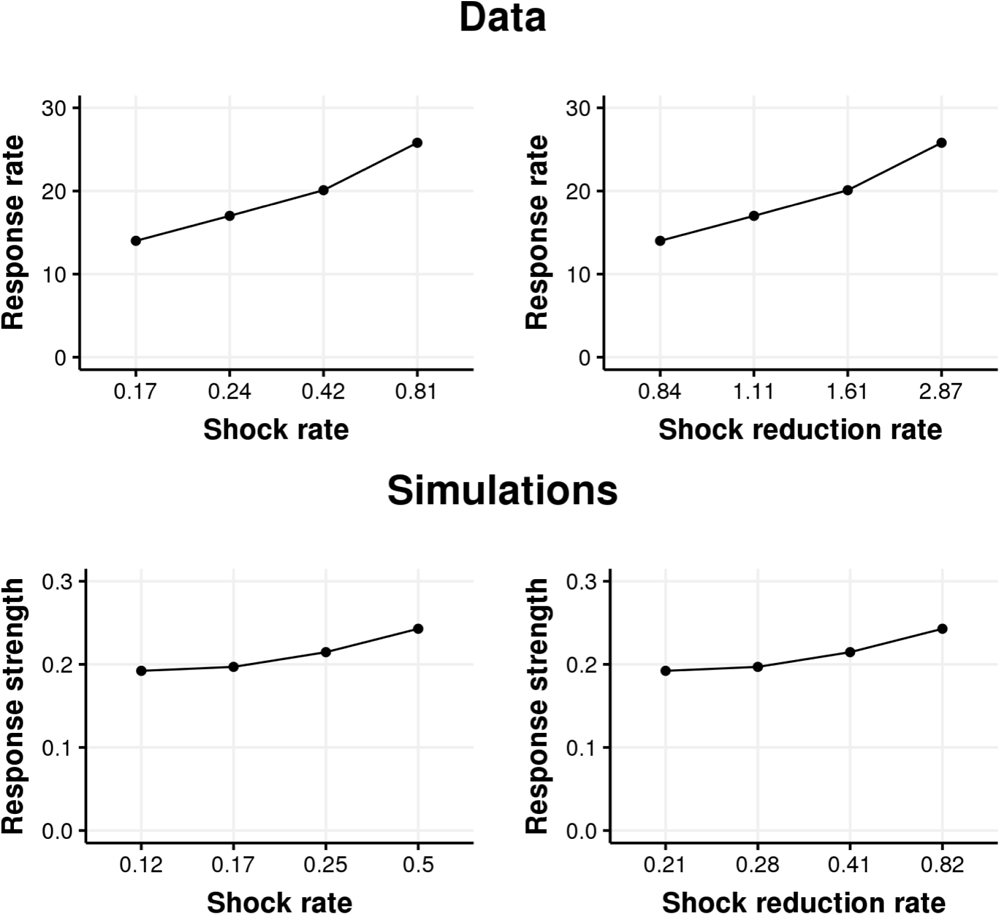
The mean responses per minute (top panels) reported by de Villiers (1974) and simulation of response strength. (*Ig*) **(bottom panels) as a function of experienced shock rate and shock rate reduction (shocks per minute) .**

### Experienced Shock Rate

More recently, Baum (2020) has concluded that the experienced shock rate is an important determinant of free-operant avoidance as a result of an extensive analysis of the relevant literature. His analysis of the de Villiers’ (1974) data showed that not only did the avoidance rate increase with shock rate reduction but also with the experienced shock rate, a relationship that is also captured by our simulations (Figure 4, left-hand panels).

In summary, simulations of goal-directed avoidance by the rate correlation mnemonic model reproduce the observed impact of the schedule shock rate, the shock rate reduction, and the experienced shock rate under two different avoidance contingencies.

## Discussion

In response to the results of Fernando and colleagues suggesting that free-operant avoidance can be conjointly controlled by goal-directed (Fernando et al., 2014a) and habitual learning (Fernando et al., 2014b), we investigated whether our dual-system model of positively reinforced free-operant behavior (Perez & Dickinson, 2020) can be extended to the corresponding negative reinforcement. Within this model, goal-directed control is determined by the experienced correlation between the response and reinforcement rates as assessed by a STM system, which in the case of avoidance yields a negative correlation. Consequently, the causal relationship between the instrumental action and the reinforcer is represented as a negative value, which interacts with the negative incentive value of the aversive reinforcer to generate a positive response strength. By simulation we demonstrated that this rate-correlation system yields avoidance rates that are positively related to two empirical correlates of free-operant avoidance, the experienced reinforcer rate and the reduction from the scheduled reinforcement rate produced by responding.

There is a dearth of theoretical accounts of free-operant avoidance. Recently, however, Baum (2020) discussed the relationship between free-operant avoidance and the experienced reinforcement or shock rate. Central to his account is the process of induction whereby experiences of a covariation, either positive or negative, between an action, such as lever pressing, and a reinforcer enables the presentations of the reinforcer to induce the action with the rate of induction being determined by the reinforcement rate. It is this process of induction that explains why the rate of lever pressing increased with the shock rate in the extensive data set considered by Baum (2020), including the de Villiers’ (1974) experiment that we addressed in our analyses. Importantly, however, the induction process is constrained by the avoidance schedule with the equilibrium response rate occurring at the point where the induction and schedule feedback functions intersect. If the response rate falls below this point the resulting increase in shock rate induces more responding, whereas if the response rate increases, the induction of responding is reduced by the decrease in shock rate. At issue in this functional account is the process by which experience of an response-reinforcer covariation produces induction. One possibility is that sensitivity to the experienced rate correlation between responding and reinforcement, which was originally identified by Baum (1973) fifty years ago as an important variable in free-operant responding, provides a process for induction, a process that is instantiated in our goal-directed system by the recycling STM mechanism.

Whatever the relationship between Baum’s induction and our goal-directed system, our dual-system model also includes a role for a second, habit system that is a version of the classic two-process account of learning in the sense that this system involves the interaction of Pavlovian conditioning and instrumental habit learning. The relevant version of the two-process account of avoidance learning is the safety signal theory of Konorski and Miller (1936), who proposed that the avoidance response is positively reinforced by its feedback stimuli through their acquisition of conditioned aversive inhibition. As in case of rewarded responding, we assume that the two systems conjointly determine the resultant response rate.

Currently, there is insufficient data to suggest how such an avoidance habit system should operate computationally to generate habit strength. In the case of rewarded or positively reinforced behavior our dual-system theory (Perez & Dickinson, 2020) proposed that the habit system employed a standard prediction-error learning algorithm that, when applied to habit learning reinforced by a safety signal, should reflect the difference between the strength of Pavlovian inhibition elicited by the feedback stimuli at the time when a response is performed and the current habit strength. In the absence of an agreed theory of Pavlovian inhibitory learning (Sosa, 2022), however, the development of a computational theory of habitual avoidance learning would be highly arbitrary.

A notable feature of rewarded or positively reinforced free-operant behavior is that the goal-directed control evident after initial training often disappears after more extended training and becomes predominantly habitual (e.g., Adams, 1982; Dickinson, Balleine, Watt, Gonzalez, & Boakes, 1995; Holland, 2004; Killcross & Coutureau, 2003). According to our dual-system theory, the loss of goal-directed control is not an inevitable consequence of extended training but depends upon whether such experience decreases the variance in response rate so that the experienced local rate correlation approaches zero. In the case of rewarded behavior which was the focus of our original theory (Perez & Dickinson, 2020), a continuously reinforced habit system yielded increasing response rates which implied a decreasing response rate variance. Whether extended training on a free-operant avoidance contingency, such as the VC schedule, reduces goal-directed control remains unknown, although Cain has discussed the neurobiological evidence indicating that avoidance transitions from goal-directed to habitual control (Cain, 2019). It should be noted, however, that responding in the Fernando et al’s (2014a) study remained sensitive to shock revaluation after 30 h of training.

A further issue concerns the motivation of free-operant avoidance that is evident in demonstrations of Pavlovian-instrumental transfer (PIT). Having trained dogs to shuttle back and forth over a barrier to avoid a shock on a free-operant avoidance schedule, Rescorla and LoLordo confined each dog on one side of the barrier where they received unavoidable shocks, each signaled by a tone. When subsequently presented while the dogs were engaged in shuttling, the tone increased the rate of responding (Rescorla & Lolordo, 1965). It is unlikely that increase was due an expectation of a shock during the tone because LoLordo subsequently demonstrated that a signal for a loud aversive klaxon produced as great an elevation of shock-reinforced avoidance responding by the dogs (LoLordo, 1967), a finding recently replicated with rats (Campese et al., 2020). Rather it would appear that the avoidance PIT reflects a general motivating effect of aversive signals.

In our discussion of positively reinforced free-operant behavior (Perez & Dickinson, 2020), we attributed general appetitive PIT to a motivational influence on habitual responding, and it is possible that general aversive PIT also operates through the habit system. To reiterate, we appealed to a Konorskian two-process mechanism whereby the habitual avoidance response was self-reinforced by the aversive Pavlovian inhibition conditioned to its feedback stimuli through their negative temporal correlation with the shock. There is evidence that this form of conditioned inhibition is mediated by the aversive excitation conditioned to the contextual stimuli. For example, Miller and colleagues reported that extinguishing the aversive excitation elicited by the contextual stimuli following inhibitory conditioning reduced the subsequent inhibition exerted by the conditioned stimuli that had been previously trained under a negative correlation with the shock in the context (Miller, Hallam, Hong, & Dufore, 1991). In this sense, inhibition is a ‘slave’ process to excitation and consequently the self-reinforcement of responding during a strong conditioned excitor will be enhanced in the PIT procedure.

It is also possible that the aversive PIT is mediated by the goal-directed system. We argued that incentive values of outcomes in the goal-directed system (Perez & Dickinson, 2020) are acquired through a process of instrumental incentive learning that enables a motivational state, such as that induced by a nutritional deficit, to control the current incentive value of a food outcome (Dickinson & Balleine, 1994). Presumably, avoidance learning, especially during the early stages, takes place while the animal is fearful with the result that this state comes to control the negative incentive value assigned to the animal’s representation of the shock, for example. As a consequence, the shock representation has a more negative incentive value when the animal is fearful than when it is not. Aversive PIT could therefore be due to an increment in negative incentive value of the shock representation during the fear-inducing signal and thereby enhance goal-directed avoidance.

Whatever the processes by which Pavlovian conditioning modulates avoidance, it is likely that it also contributes to the extinction of avoidance through the extinction of aversive excitatory conditioning to the context. According to our dual-system account (Perez & Dickinson, 2020) extinction of rewarded behavior is a complex of interacting systems. As the rate correlation is undefined when the STM is cleared of any reinforcer representations following the onset of extinction, *g* remains fixed at the weighted average value after the last recycle containing an outcome representation. However, as *g* predicts the same rewarding outcome that reinforced the habit, *g* contributes to the prediction error generating the habit strength, *h*. As a consequence, during extinction *h* acquires a sufficient negative value to counteract the contribution of the terminal positive *g* to responding. By contrast, this interaction does not occur in avoidance because the outcome represented by the goal-directed system and the reinforcer of habits are different events: the shock and the feedback stimuli, respectively. Therefore, the residual *g* could maintain persistent responding in extinction.

This predicted persistence might be thought to be a virtue of the model as it is often claimed that avoidance responding is abnormally persistent in extinction. However, there is little reason to believe that persistence is a feature of free-operant avoidance. Perhaps the most pertinent study is one by Uhl and Eichbauer in which rats were trained to avoid a shock by lever pressing on a VC schedule similar to that employed by de Villiers (1974) before responding was extinguished by omitting the shock (Uhl & Eichbauer, 1975). Persistence of avoidance responding was contrasted with that following positively reinforced VC training in which a lever press during a cycle delivered sugar water to hungry rats at the end of the cycle. Following both positive (reward) and negative (avoidance) reinforcement training, extinction was similarly rapid with responding during the first 3-h extinction session being about a fifth of that at the end of training which Uhl and Eichbauer attributed to generalization decrement produced by the absence of the reinforcer (Uhl & Eichbauer, 1975). In agreement with this account, transferring from extinction to a non-contingent shock presentations under a variable time (VT) schedule immediately reinstated avoidance responding to the level seen in the last reinforced session.

This marked loss of performance in extinction contrasted with that observed when the rats were transferred from the VC to a VT schedule under which the reinforcer is delivered with the same recycling time as during training but independently of responding. Although relative to the terminal VC rates, initial avoidance was marginally more persistent than rewarded responding, in both cases the level was greatly elevated above that in extinction before progressively declining to a low level. Contrasting performance under the VC schedule with that under the VT schedule rather than standard extinction provides an unconfounded measure of the impact of positive (reward) and negative (avoidance) instrumental contingencies by controlling the role of the discriminative and Pavlovian functions of reinforcement in producing generalization decrements. For both rewarded and avoidance lever pressing by rats, the goal-directed system predicts a progressive decline of responding as *g* gradually decreases with the accumulation of STM recycles with a zero rate rate correlation. Similarly, the feedback stimuli lose their inhibitory properties as there is no longer a positive Pavlovian contingency between these stimuli and the reinforcer. In summary, in accord with Uhl and Eichbauer’s (1975) results, our dual-system theory predicts positive and negative reinforcement training should yield similar profiles of responding under a non-contingent schedule as a common rate correlation mechanism mediates rewarded responding and avoidance.

We have already noted that the STM model of the rate correlation system could well yield a non-zero value for *g*, positive for reward training and negative for avoidance, after extinction. This feature of the model explains Rescorla’s finding that sensitivity to outcome revaluation of a positive reinforcer persists after extinction (Rescorla, 1993). In contrast, the model predicts that exposure to non-contingent positive reinforcement should reduce goal-directed control, a prediction recently confirmed by Crimmins and colleagues once Pavlovian influences are controlled (Crimmins, McNulty, Laurent, Hart, & Balleine, 2022). Our rate correlation makes exactly the same predictions in the case of free-operant avoidance. Revaluing the aversive reinforcer, for example, by presentations of the shock under morphine (Fernando et al, 2014a) should reduce subsequent avoidance following extinction, but not following non-contingent shock exposure.

When taken in conjunction with Perez and Dickinson (2020), the dual system model of free-operant behavior that incorporates a rate correlation system accounts for two of the four interactions between the response-reinforcer contingency and the valence of the reinforcer: positive reinforcement (reward) when both these factors are positive and negative reinforcement (avoidance) when both factors are negative. The role of goal-directed control in the remaining two interactions, omission and punishment, remains to be determined. In the case of an omission schedule under which the contingency is negative and valence positive, Dickinson and colleagues (1998) failed to find any evidence for goal-directed control: devaluing the omitted reinforcer had no impact on the level of response reduction produced by prior omission training.

To the best of our knowledge, whether or not the response reduction produced by free-operant punishment, under which the contingency is positive and the outcome valence is negative, has a goal-directed component has not been investigated. Rate correlation theory anticipates that the negative *Ig* product generated by the punishment will detract from the positive *Ig* product for the reward contingency used to establish responding. Given this analysis, the rate correlation account of goal-directed responding predicts that decreasing the negativity of the incentive value of the punisher by outcome revaluation should immediately reduce the response suppression previously established by punishment.

## Acknowledgments

This research was partially funded by ANID-SIA 85220023 and ANID FONDECYT 1231027, awarded to Omar D. Perez, and Instituto Sistemas Complejos de Ingeniería ANID-PIA/PUENTE AFB220003.

## Footnotes

1 The feedback stimulus was also omitted during this test to ensure that revaluation of shock did not act through the reinforcing properties of the stimulus rather than directly through a change in the value of the shock itself. Of course, this procedure did not eliminate any effect mediated by the intrinsic feedback stimuli.

2 This procedure for deleting memory samples differs from that used by Perez and Dickinson (2020), which deleted the oldest sample at a recycle. The original procedure requires the system to represent the age of a sample, whereas random deletion ensures that the older a sample the more likely it is to have been deleted at any given recycle without requiring a representation of its age. Using the Perez and Dickinson (2020)’s procedure did not change the results of the simulations.

3 In their paper, the authors employed probabilities of shock per 2 seconds. Here we work with equivalent probabilities per 1 second. The reason for this choice is that 1-second timebase is the one that we have employed for simulations of free-operant responding under positive reinforcement (Perez & Dickinson, 2020) and the acquisition of avoidance on the simple VC (see Figure 2).

## Notes

### Competing Interest Statement

The authors have declared no competing interest.

### Summary of Updates

Responded to reviewers and added new simulations.

